# *Pseudomonas syringae* effector AvrE associates with plant membrane nanodomains and binds phosphatidylinositides i*n vitro*

**DOI:** 10.1101/2021.07.08.451616

**Authors:** Xiu-Fang Xin, Lisa Kinch, Boying Cai, Bradley C. Paasch, Brian Kvitko, Nick V. Grishin, Sheng Yang He

## Abstract

Bacterial phytopathogens deliver effector proteins into host cells as key virulence weapons to cause disease. Extensive studies revealed diverse functions and biochemical properties of different effector proteins from pathogens. In this study, we show that the *Pseudomonas syringae* effector AvrE, the founding member of a broadly conserved and pathologically important bacterial effector family, binds to phosphatidylinositides (PIPs) *in vitro* and shares some properties with eukaryotic PROPPINs (β-propellers that bind polyphosphoinositides). In planta pull down experiments with transgenic Arabidopsis plants expressing AvrE revealed that AvrE is associated with several plant proteins including plasma membrane lipid-raft proteins. These results shed new light on the properties of a bacterial effector that is crucial for bacterial virulence in plants.

## INTRODUCTION

Many Gram-negative pathogenic bacteria of animals and plants utilize a type III secretion system (T3SS) to deliver effector proteins into the host cell as a common mechanism of pathogenesis (Galan and Collmer, 1999; He et al., 2004). Among the most crucial virulence factors of plant pathogenic bacteria is the AvrE family of effectors, of which AvrE from *Pseudomonas syringae* pv. *tomato* (*Pst*) strain DC3000 is the founding member (Lorang and Keen, 1995). The AvrE effector family is present in diverse plant pathogenic bacteria that belong to the genera of *Pseudomonas*, *Pantoea*, *Erwinia*, *Dickeya*, and *Pectobacterium*. In *Pst* DC3000, AvrE is functionally redundant to another effector, HopM1 (Alfano et al., 2000; Badel et al., 2006; DebRoy et al., 2004). Mutation of either *avrE* or *hopM1* alone does not strongly affect *Pst* DC3000 virulence, but the *avrE hopM1* double mutant is severely impaired in virulence (Badel et al., 2006). In bacteria that lack the *hopM1* gene, mutation of *avrE* orthologues alone has been shown to cause a dramatic decrease in virulence(Bogdanove et al., 1998; Boureau et al., 2006; DebRoy et al., 2004; Frederick et al., 2001; Gaudriault et al., 1997). Because of the important role of AvrE-family effectors in bacteirla virulence and the wide distribution of this effector family in diverse bacteria, an understanding of the properties of AvrE-family effectors is important for understanding bacterial pathogenesis in plants.

AvrE-family effectors, including AvrE from *Pst* DC3000, WtsE from *P. stewartii*, DspE from *E. amylovora*, and DspE from *Pectobacterium carotovorum*, have been shown to cause water-soaking and/or cell death when expressed in host or non-host plants (Boureau et al., 2006; Degrave et al., 2008; Frederick et al., 2001; Ham et al., 2008; Ham et al., 2006; Ham et al., 2009; Hogan et al., 2013; Kim et al., 2011; Xin et al., 2015) and can suppress plant defense responses such as callose deposition and immune-related gene expression (Boureau et al., 2006; DebRoy et al., 2004; Ham et al., 2008; Ham et al., 2009; Jin et al., 2016; Xin et al., 2015). DspE from *E. amylovora* was reported to interact with several putative receptor kinases from apple (Meng et al., 2006), whereas WtsE from *P. stewartii* has been shown to target protein phosphatases PP2A B’ subunits in plants (Jin et al., 2016). The mechanisms by which DspE and WtsE induce water soaking and cell death or suppress defense responses via targeting plant receptor kinases and PP2As are not understood.

In a previous study, we found that AvrE is localized to the plasma membrane (PM) in a punctate pattern (Xin et al., 2015). In the current study, we found that AvrE binds to phosphatidylinositides (PIPs) and shares some features with the eukaryotic PROPPIN (β-propellers that bind polyphosphoinositides) family of proteins (Baskaran et al., 2012; Dove et al., 2004; Krick et al., 2012). In addition, we found that AvrE is associated with several PM lipid-raft proteins *in vivo*, suggesting that AvrE likely exerts its virulence actions, including modulation of water-soaking, cell death and defense responses, within PM lipid rafts in plants.

## RESULTS

### AvrE is associated with PM “lipid raft/membrane microdomain” proteins in Arabidopsis

To determine whether AvrE is localized in the PM in association with specific host proteins, we carried out *in vivo* protein pull down experiments using total protein extracts from His:YFP:AvrE transgenic plants. The AvrE expression level was optimized in DEX:*His:YFP:AvrE* Arabidopsis plants so that enough AvrE protein could be expressed in leaves without causing too quick tissue collapse. Protein pull-down experiments were conducted using GFP-Trap-A beads (see MATERIALS and METHODS), and Arabidopsis proteins pulled down by His:YFP:AvrE were analyzed and identified by mass spectrometry (Table 1). Interestingly, the majority of Arabidopsis proteins pulled down by AvrE were either known or predicted PM proteins. Moreover, many of the proteins identified, including Hypersensitive Induced Reaction Protein 2 (HIR2), Plasma membrane Intrinsic Proteins (PIPs) and Arabidopsis H^+^-ATPases (AHAs), are classical “lipid rafts/membrane microdomain” proteins in the PM (Borner et al., 2005; Keinath et al., 2010; Mongrand et al., 2004; Shahollari, 2004). Taken together, these results are consistent with the previous finding that AvrE is targeted to the host PM (Xin et al., 2015), and, moreover, suggest that AvrE is localized within the lipid rafts/membrane microdomains in the PM.

**Table 1.** Arabidopsis proteins pulled down by AvrE.

| **Gene Annotation** | **Gene Identifier** | **Peptide count** | **Times** | **PM/Lipid raft localization** |
| --- | --- | --- | --- | --- |
| AvrE |  | 206 | 3 |  |
| HIR2; SPFH/Band 7/PHB domain-containing membrane-associated protein family | AT3G01290 | 13 | 3 | Yes<sup>a</sup> |
| PIP1B; plasma membrane intrinsic protein 1B | AT2G45960 | 4 | 2 | Yes<sup>a</sup> |
| PIP2A; plasma membrane intrinsic protein 2A | AT3G53420 | 11 | 3 | Yes<sup>a</sup> |
| PIP3A; plasma membrane intrinsic protein 3 | AT4G35100 | 6 | 2 | Yes<sup>a</sup> |
| AHA1; H(+)-ATPase 1 | AT2G18960 | 50 | 3 | Yes<sup>a</sup> |
| ERD4; Early-responsive to dehydration stress protein (ERD4) | AT1G30360 | 11 | 2 | Yes<sup>a</sup> |
| PEN3; Plant PDR ABC-type transporter family protein | AT1G59870 | 30 | 2 | Yes<sup>b</sup> |
| TOPP1; type one serine/threonine protein phosphatase 1 | AT2G29400 | 5 | 3 |  |
| TOPP2; type one serine/threonine protein phosphatase 2 | AT5G59160 | 33 | 3 |  |
| TOPP5; type one protein phosphatase 5 | AT3G46820 | 7 | 3 |  |
| CPK7; calmodulin-domain protein kinase 7 | AT5G12480 | 7 | 2 | Yes<sup>b</sup> |
| CPK32; calcium-dependent protein kinase 32 | AT3G57530 | 14 | 3 | Yes<sup>b</sup> |

A list of proteins pulled down by His:YFP:AvrE as bait. Three repeats were performed with similar results, and proteins pulled down in at least two repeats are shown. “Peptide count” for each protein from the 2<sup>nd</sup> repeat is presented. In the “PM/lipid raft localization” column, “Yes” indicates the protein is known to be localized in the plant PM. <sup>a</sup>, proteins previously found to be enriched in lipid rafts (biochemically isolated Detergent Resistant Membrane fraction from plant extract) (Borner et al., 2005; Keinath et al., 2010; Kierszniowska et al., 2009; Shahollari, 2004). <sup>b</sup>, proteins in the same gene family were previously identified as enriched in lipid rafts.

### The N-terminal half of AvrE is predicted to form a β-propeller and binds to phosphatidylinositides

Next, we conducted bioinformatic analysis of the AvrE sequence using PSI-BLAST and HHPRED (Altschul et al., 1997; Soding et al., 2005), which revealed that AvrE contains a β-propeller in the N-terminal half of the protein (probability as high as 97.4). While the top HHPRED hit of AvrE matches a 7-bladed propeller, many of the lower hits mapped AvrE to a duplicated β-propeller with similar probabilities (Table 2). Given the longer than usual sequence range containing WD40 repeats (523 residues or 13 repeating units), the AvrE N-terminus may adopt a duplicated β-propeller fold. A similar β-propeller domain was also predicted for *E. amylovora* DspE, which shares only 23% amino acid identity with AvrE (Sabrina et al., 2013), suggesting conservation of this domain in divergent members of AvrE-family effectors.

**Table 2.**
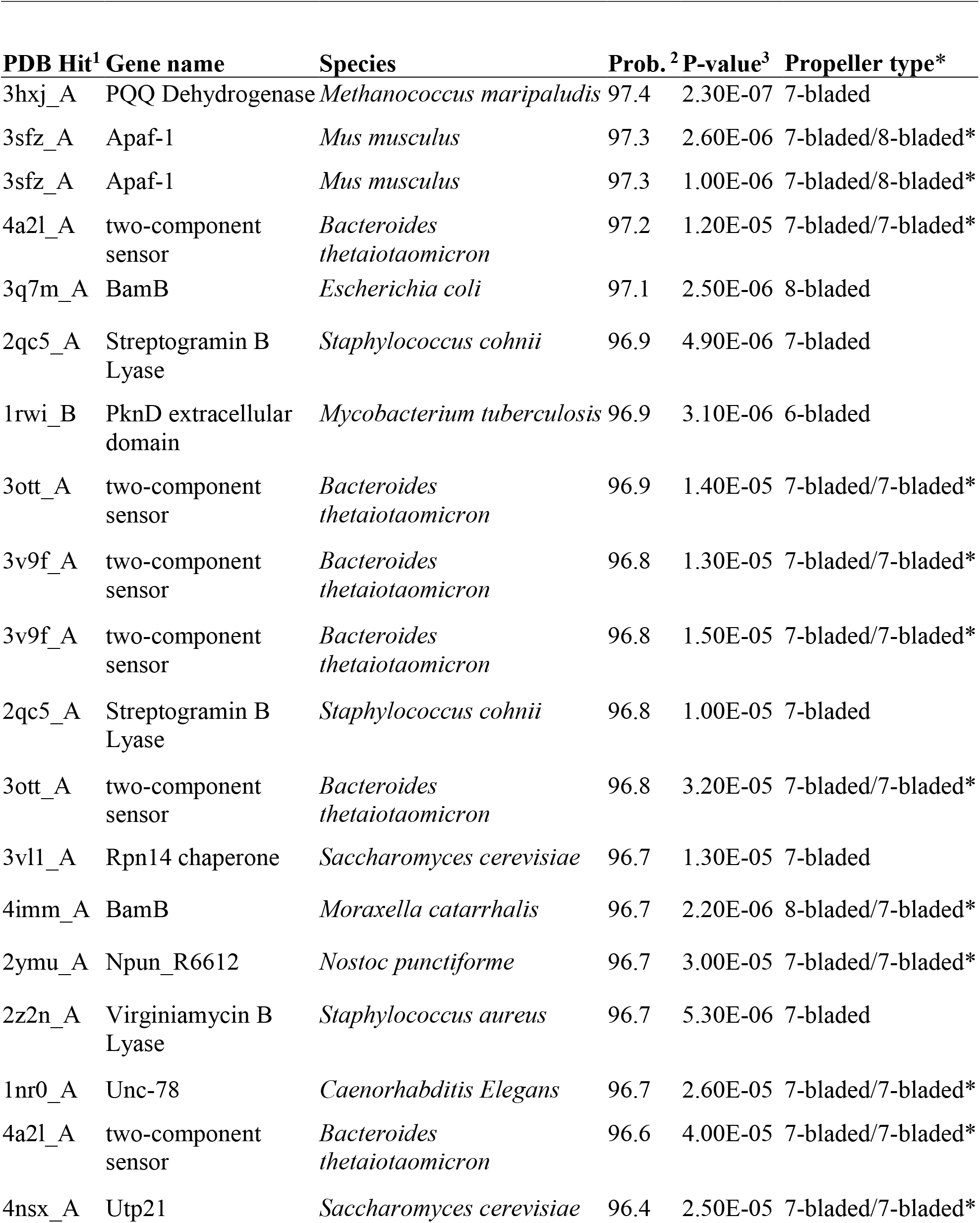
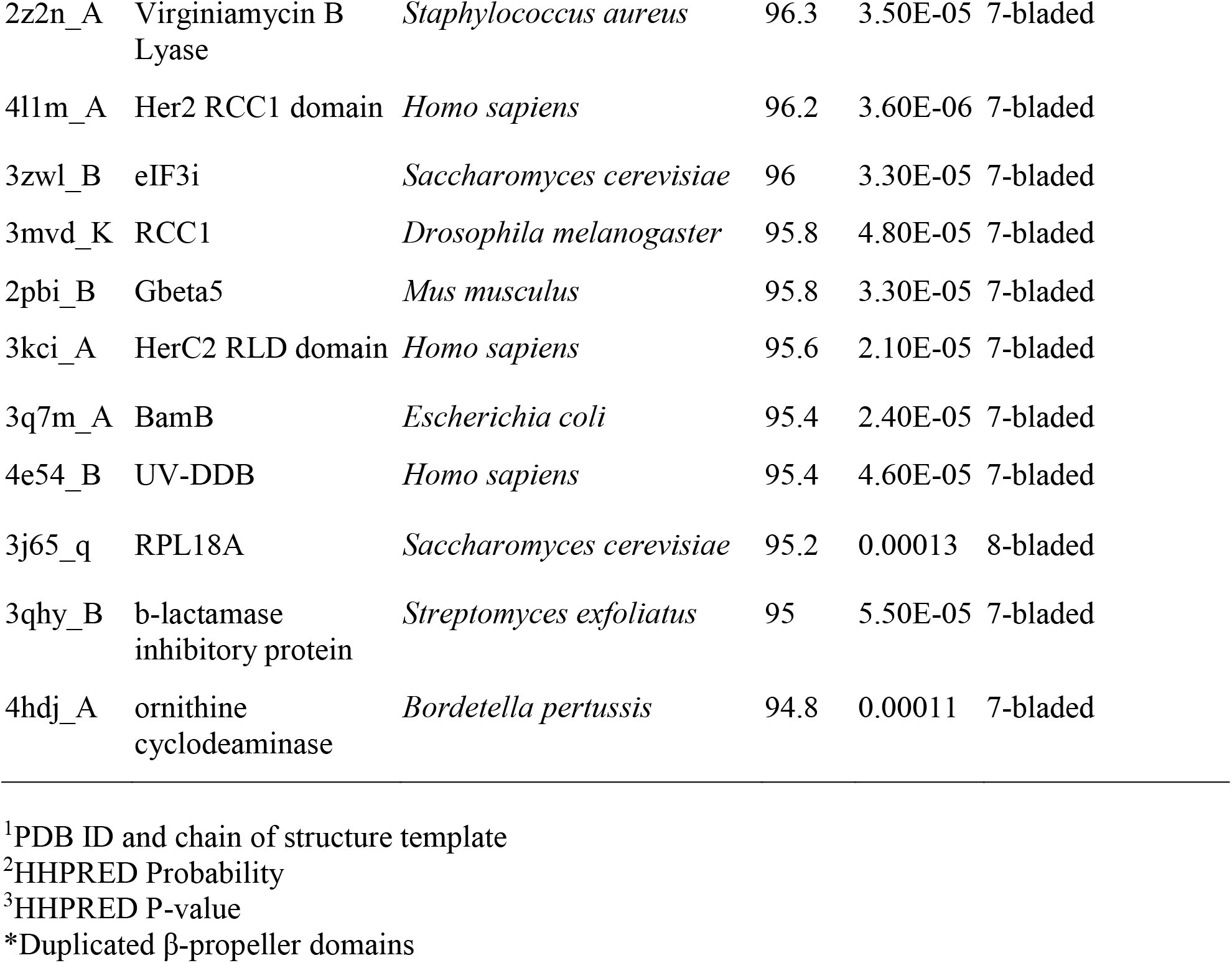
Top30 HHPRED Structures identified by AvrE1 N-terminus.

A specific family of eukaryotic 7-bladed β-propeller proteins binds to phosphatidylinositol phosphates, PI3P, PI(3,5)P2, and PI(3,4,5)P_3_ (Baskaran et al., 2012; Dove et al., 2004; Krick et al., 2012). These proteins are called PROPPINs (β-propellers that bind polyphosphoinositides) (Dove et al., 2009; Dove et al., 2004). We tested the possibility that AvrE might bind to PIPs by incubating purified MBP-AvrE-His with a PIP lipid strip (P-6001; Echelon Biosciences). Remarkably, at the concentration as low as 1 nM, AvrE binds to three PIPs, PI(3)P, PI(3,5)P2 and PI(3,4,5)P3, but not other PIPs, phosphatidic acid (PA), phosphatidylinositol (PI), phosphatidylethanolamine (PE), phosphatidylchorine (PC), phosphatidylserine (PS) or sphingosine-1-phosphate (S1P) (Fig. 1A, B). To determine whether the β-propeller region of AvrE binds to PIPs, we purified the N-terminal 700-aa peptide (AvrE_1-700_). The C-terminal 595-aa peptide (AvrE_1201-1795_) was used as a control. As shown in Fig. 1B, AvrE_1-700_ also binds to PI(3,5)P2, PI(3,4,5)P3 and, to less extent, PI(3)P, whereas AvrE_1201-1975_ had no PIP-binding activity. This result demonstrates that a PIP-binding motif is localized within AvrE_1-700_. Since PI(3,4,5)P3 has not been found in plants, we performed liposome binding assay to confirm the interaction between purified MBP-AvrE-His protein and PI(3)P and PI(3,5)P2 (Fig. 1C).

**Figure 1.**
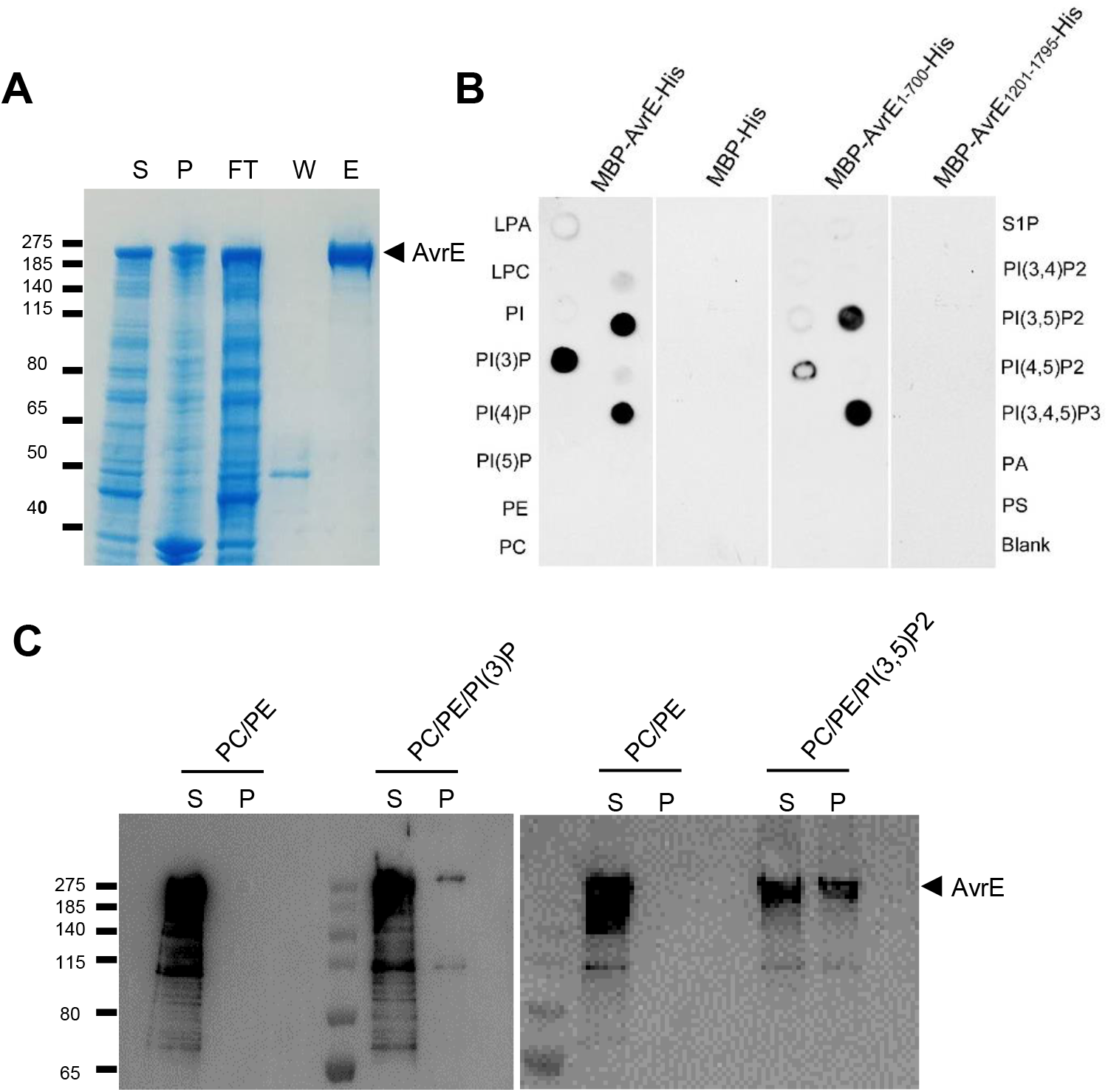
AvrE protein binds to PIPs *in vitro*. (**A**) Purification of MBP-AvrE-His from *E. coli* cells. S, supernatant; P, pellet; FT, flow through (after affinity purification); W, wash through; E, elution. ((**B**) Full-length AvrE and AvrE_1-700_ binds to PIPs. Lipid overlay assays of purified MBP-AvrE (1 nM), MBP (10 nM), MBP-AvrE_1-700_ (10 nM) and MBP-AvrE_1201-1795_ (10 nM). PIPs: PI(3)P, PI(3,5)P2 and PI(3,4,5)P3, phosphatidic acid (PA), phosphatidylinositol (PI), phosphatidylethanolamine (PE), phosphatidylchorine (PC), phosphatidylserine (PS) or sphingosine-1-phosphate (S1P), lysophosphatidic acid (LPA), lysophosphatidylchorine (LPC). (**C**) Liposome binding assay. Liposomes were incubated with purified AvrE protein (from **A**). After ultracentrifugation, the liposome-free supernatant (S) and liposome pellet (P) were analyzed by SDS–PAGE followed by anti-AvrE-N immunoblot.

Eukaryotic PROPPINs contain a conserved “FRRG” motif in which the arginine residues play a key role in binding to PIPs (Baskaran et al., 2012; Krick et al., 2012). However, our bioinformatic analysis did not reveal such a motif in AvrE, suggesting that AvrE has evolved an alternative PIP-binding motif. Through an alignment of AvrE-family effector proteins, we found two highly conserved “GxxY” motifs in the predicted β-propeller domain of AvrE (Fig. 2A). In the predicted β-propeller structure of AvrE (using the top hit of 7-bladed β-propeller protein as a template), each GxxY motif forms a conserved part of a “propeller blade” and marks a turn that starts the second β-strand of a blade (Fig. 2A). Because the GxxY motif does not contain arginine residues, it is not likely to be responsible for PIP-binding. Nevertheless, we determined whether the highly conserved GxxY motif is important for the virulence function of AvrE. The conserved tyrosine (Y) residue was mutated to alanine (A) in the two GxxY motifs (i.e., Y416A, Y510A), and the mutant proteins were transiently expressed in *N. benthamiana.* While single Y-to-A mutations in either motif (AvrE-Y416A, hereinafter AvrE-y1; AvrE-Y510A, hereinafter AvrE-y2) did not affect the ability of AvrE to induce tissue necrosis, double mutations (i.e., AvrE-yy) completely abolished it (Fig. 2B). We then tested the effect of these mutations on AvrE function during *Pst* DC3000 infection of *N. benthamiana*. Plasmids carrying wild-type (pBBR1:*avrF:avrE:HA*; pCPP5961 (Kvitko et al., 2009)) or mutant *avrE* (pBBR1:*avrF:avrE*-yy*:HA*) genes were constructed and introduced into the ΔCEL mutant (*avrE^−^, hopM1^−^*) in the Δ*hopQ1* mutant background of *Pst* DC3000 (Kvitko et al., 2009), which multiplies in *N. benthamiana* plants (Wei et al., 2007). Translocation assay showed that AvrE-yy protein was normally translocated into the plant cell (Fig. S1). However, whereas *avrE*-WT completely complemented the virulence defect of the Δ*hopQ1*ΔCEL mutant in *N. benthamiana*, the *avrE*-yy mutant could not (Fig. 2C), demonstrating the critical importance of the GxxY motif for the virulence function of AvrE.

**Figure 2.**
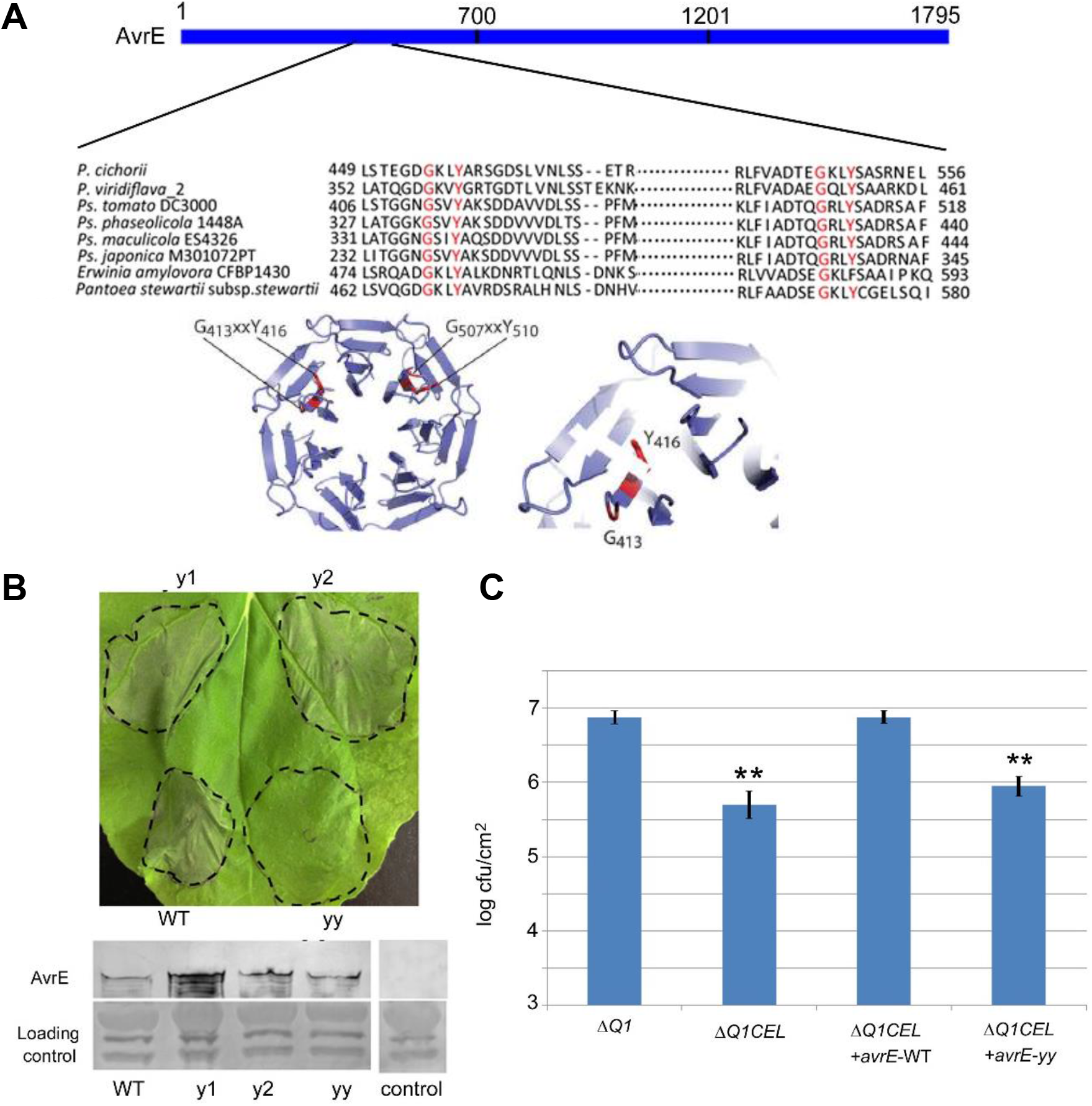
AvrE is predicted to form a β-propeller and a “GxxY” motif is essential for its virulence function. (**A**) A diagram showing the two GxxY motifs (upper) and the predicted β-propeller structure at the N terminus of AvrE (lower). Note that AvrE may form a double β-propeller and here a 7-bladed β-propeller based on one of the top HHPRED hits (2qc5) was shown on the left, with the relative amino acid positions (413-416; 507-510) of GxxY motifs indicated. A zoom of one GxxY motif shown on the right. **(B)** The double Y to A mutations (yy) in the two GxxY motifs abolished the ability of AvrE to cause tissue necrosis. *Agrobacterium* containing β-estradiol:*avrE* WT:VFP, y1:VFP, y2:VFP, yy:VFP plasmids were infiltrated into the leaves of *N. benthamiana* and 20 μM β-estradiol were sprayed 24 hours after *Agrobacterium* infiltration. Pictures were taken 2 days after β-estradiol treatment. The lower part shows AvrE proteins detected in western blot for all the constructs. A rabbit poly-clonal AvrE-N antibody was used. **(C)** The *avrE*-yy mutant failed to complement the growth of Δ*hopQ1*ΔCEL mutant in *N. benthamiana*. Δ*Q*1: Δ*hopQ1*mutant of *Pst* DC3000; Δ*Q1*CEL: Δ*hopQ1*ΔCEL mutant of *Pst* DC3000; avrE-WT: pBBR1:*avrF:avrE*:*HA*; avrE-yy: pBBR1:*avrF:avrE*-yy:*HA* > ** indicates a significant difference between to the Δ*Q*1 mutant and the rest of strains; P value <0.01.

## DISCUSSION

Lipid rafts/microdomains are defined as small (10-200nm) and highly dynamic, sterol- and sphingolipid-enriched domains on the PM. They are believed to be a compartmentalization mechanism that enables cells to carry out specialized cellular functions in response to environmental stimuli (Cacas et al., 2012; Malinsky et al., 2013). Furthermore, lipid rafts are enriched for polyphosphoinositides (Furt et al., 2010). The finding of AvrE association with plant lipid raft proteins warrants discussion. First, the punctuate/vesicle-like localization pattern of AvrE on the plant PM (Xin et al., 2015) is consistent with the lipid raft/microdomain association in plants, because this localization pattern is reminiscent of that of several known plant lipid raft/microdomain proteins (Li et al., 2012; Qi et al., 2011; Varet et al., 2003). In addition, a previous study showed that *E. amylovora* DspA/E (an orthologue of AvrE) had a negative effect on the accumulation of certain sphingolipid biosynthetic intermediates in yeast cells (Siamer et al., 2014). Although it is not known whether DspA/E or AvrE has this effect in plant cells, this intriguing observation is consistent with the localization of AvrE in sphingolipid-rich lipid rafts observed in this study. Second, there is emerging evidence showing that plant lipid raft may be an important platform for immune signaling. For instance, studies showed that the flagellin receptor FLS2 is localized to and enriched in membrane microdomains after flg22 treatment (Bücherl et al., 2017; Keinath et al., 2010). The motility of FLS2 in the PM is decreased after PAMP elicitation (Ali et al., 2007). Further support for this was provided by the observation that FLS2 interacts with HIR2 (Qi and Katagiri, 2012), a known lipid raft-associated protein and one of the lipid raft/microdomain proteins associated with AvrE (Borner et al., 2005; Keinath et al., 2010; Liu et al., 2009) (Table 1). It is possible that AvrE may interfere with immunity-related lipid rafts in plants to disrupt immune function. However, our genome-wide microarray analysis did not reveal global effects of AvrE on defense gene expression associated with FLS2 (Xin et al., 2015). This result suggests that AvrE association with lipid raft proteins likely affects other plant processes independent of FLS2 signaling.

The discovery that AvrE shares several conserved features with eukaryotic 7-bladder β-propeller domain-containing PROPPINs was made through a combination of bioinformatics and biochemical analyses. We found that the β-propeller domain of AvrE-family effectors contains two highly conserved GxxY motifs. Site-directed mutagenesis revealed that these two motifs play a critical role in the virulence function of AvrE (Fig. 2B, C). Because the β-propeller domain is also predicted for DspE, which is distantly related to AvrE in the AvrE family (Sabrina et al., 2013), we propose that the β-propeller domain-based PIP-binding is likely a shared biochemical and structural feature of all AvrE-family effectors and that AvrE-family effectors are potential bacterial mimics of eukaryotic PROPPINs. The eukaryotic PROPPINs contain a conserved “FRRG” motif that binds to PIPs (Baskaran et al., 2012; Krick et al., 2012). However, bioinformatic analysis did not reveal such a motif in AvrE. This suggests that AvrE-family effectors may have evolved a novel PIP-binding motif in the β-propeller domain. Eukaryotic PROPPINs mediate membrane trafficking at many sites in the cell, including those essential for autophagy (Baskaran et al., 2012; Dove et al., 2009; Krick et al., 2012). Because a prominent output of the AvrE virulence activity is host cell death, it would be interesting in the future to determine whether AvrE induces host cell death via autophagy (Lal et al., 2020).

In summary, many mammalian and plant pathogenic bacteria use type III effector proteins to target host cellular processes as a central mechanism of pathogenesis. AvrE-family effector proteins are among the most important virulence factors in plant-pathogenic bacteria and are widely conserved in the divergent genera of *Pseudomonas*, *Pantoea*, *Erwinia*, *Dickeya*, and *Pectobacterium*. Notably, these effectors are encoded by genes that are physically adjacent to the T3SS gene cluster as part of a large pathogenicity island (Bogdanove et al., 1998; Boureau et al., 2006; DebRoy et al., 2004; Frederick et al., 2001; Gaudriault et al., 1997), acquisition of which is believed to be central in the evolution of many bacterial plant pathogens. As such, targeting plant PM lipid rafts via a novel type of PIP-binding effectors, as shown for AvrE in this study, may represent a fundamental and previously enigmatic aspect of bacterial pathogenesis in plants.

## MATERIALS and METHODS

### Molecular cloning

The *avrE*-yy mutation was made using the QuickChange II site-directed mutagenesis kit (Agilent Technologies). pENTR/D-TOPO:*avrE* was used as a template for quick change PCR; primers used were Y416A F/R and Y510A F/R (Table S1). The constructs of β-estradiol:*avrE*/y1/y2/yy:*VFP* were cloned by gateway cloning (Life Technologies). pBBR1:*avrF:avrE*-yy:*HA* was constructed by *Rsr*II sub-cloning of the fragment containing yy mutation from pENTR/D-TOPO: *avrE*-yy into *Rsr*II digested pBBR1:*avrF:avrE*:*HA* (pCPP5961) (Kvitko et al., 2009). pENTR/SD/D:*avrF:avrE-yy* was constructed by *Asc*I subcloning from pBBR1:*avrF:avrE*-yy:*HA* to replace the *avrE* WT allele in pENTR/SD/D: *avrF:avrE* (pCPP5912) (Kvitko et al., 2009). Cya translocation reporter constructs were created by LR reaction using pENTR/SD/D: *avrF:avrE*, pENTR/SD/D:*avrF:avrE-yy* and the destination vector pBBR1: GW:RfB:Cya (pCPP5371) (Oh et al., 2007).

#### Bioinformatic analysis and structure prediction of AvrE

PSI-BLAST(Altschul et al., 1997) initiated with the N-terminal half of AvrE (gi|28851823 residues 1-600) identified a β-propeller/WD40 repeats from various protein sequences starting in the second iteration. To further support the presence of WD40 repeats in the AvrE N-terminus and to map the sequence to structure, we queried the PDB70 database using HHPRED (Soding et al., 2005). The AvrE N-terminus confidently identified various β-propeller structures as top hits. The top-scoring structures map to various different AvrE sequence ranges, with each alignment starting and ending at the different consecutive WD40 repeats that encompass AvrE residues 309-832.

#### Protein purification from *E. coli* and lipid binding assays

The *avrE*-wild type, *avrE*-yy, *avrE_1-700_* and *avrE_1201-1795_* genes were cloned onto the vector pET-MAL-c4x (Melotto et al., 2008) at the *Bam*HI*/Xho*I restriction sites, to generate constructs expressing AvrE proteins tagged with MBP and His at the N-or C-terminus, respectively. Primers used were AvrE F/R, AvrE700R and AvrE1201F (Table S1). Expression plasmids were transformed into the Rosetta 2 strain of *E. coli* for protein induction. Soluble protein extracts were purified using amylose resin (New England Biolabs) and/or Ni-NTA agarose beads (Life Technologies).

Lipid overlay assays were performed using membrane lipid strips (P-6001; Echelon Biosciences) and purified AvrE or MBP proteins. PBS buffer + 0.1% Tween 20 + 1 mM MgCl_2_ was used as incubation and washing buffer. An anti-MBP antibody (1:5000 dilution; Sigma) was used for detecting the proteins.

For liposome binding assay, phosphatidylcholines, phosphatidylethanolamines and phosphoinositides were dissolved in “65% chloroform+35% methanol”. Lipids with indicated compositions were mixed in a glass vial. The solvent was evaporated under a stream of nitrogen in a chemical hood. Multilamellar liposomes were formed by rehydrating the lipid film in a rehydration buffer (20 mM Tris–HCl, pH 7.6, 100 mM NaCl) at 63°C for 1 h. Liposomes were extruded through a polycarbonate filter (400 nm or 100 nm) for 25 times using a mini-extruder (Avanti Polar Lipids). Purified AvrE protein was incubated with the liposome at room temperature for 1 hour. Samples were centrifuged in ultracentrifuge at 100,000g for 15 min at 4 °C. The supernatant (S) was collected to examine proteins not bound to the liposome. The pellets (P) were washed twice with 100 μl rehydration buffer. The S and P fractions were analyzed by SDS–PAGE followed by anti-AvrE-N immunoblot.

#### Bacterial disease assays

*Nicotiana benthamiana* plants were grown in a growth chamber with a 12-hour-day/12-hour-night cycle (~200 μE fluorescent light, 24 ºC). Disease assays with *N. benthamiana* plants was performed as previously described (Kvitko et al., 2009). Briefly, leaves of 8-week-old *N. benthamiana* were infiltrated with Δ*hopQ1*, Δ*hopQ1*ΔCEL, Δ*hopQ1*ΔCEL+*avrE-*WT:HA, or Δ*hopQ1*ΔCEL+*avrE*-yy:HA at 1×10^4^cfu/ml. Bacteria populations were determined at day 4 post-inoculation.

#### Cya calmodulin-dependent adenylate cyclase effector translocation assay

Leaves of 6-week-old *N. benthamiana* were infiltrated with Δ*hopQ1*ΔCEL+pBBR1*:avrF:avrE-WT:Cya*, Δ*hopQ1*ΔCEL+ pBBR1:*avrF:avrE-yy:Cya* or Δ*hopQ1 hrcC+*pBBR1:*avrF:avrE-yy:Cya* at 5×10^7^ cfu/ml. Tissue samples from infiltrated areas were collected 7 h post infiltration and frozen in liquid N_2_. The concentration of cAMP in infiltrated tissue were determined using the Direct cAMP ELISA kit (Enzo Life Science) and the concentrations of protein were determined using Quick Bradford Assay Solution (Bio-Rad).

## ACKNOWLEDGEMENTS

We thank Kim Orth and Dor Salomon for the help with bioinformatics analysis of AvrE. This work was supported by funding to S.Y.H. from the National Institutes of Health R01AI060761 and R01 AI0155441 and to X.F.X. from National Natural Science Foundation of China (Grant number 31870233) and Chinese Academy of Sciences Strategic Priority Research Program (Type-B; Grant number XDB27040211).

## AUTHOR CONTRIBUTIONS

Conceived and designed the experiments: XFX SYH. Performed the experiments: XFX LK BC BCP BK. Wrote the paper: XFX SYH, with input from all authors.

---

**Table S1.**
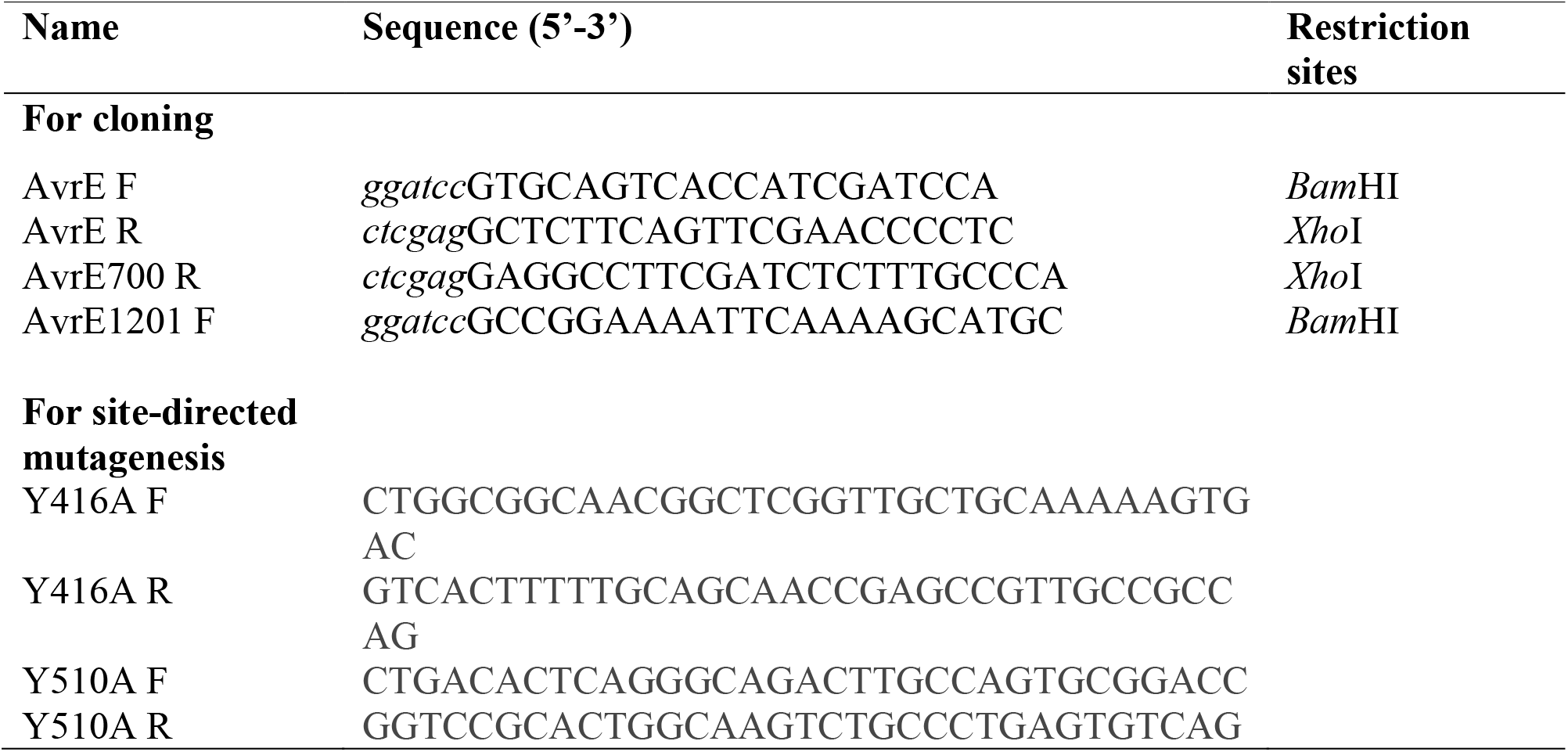
Primers used in this study.

**Figure S1.**
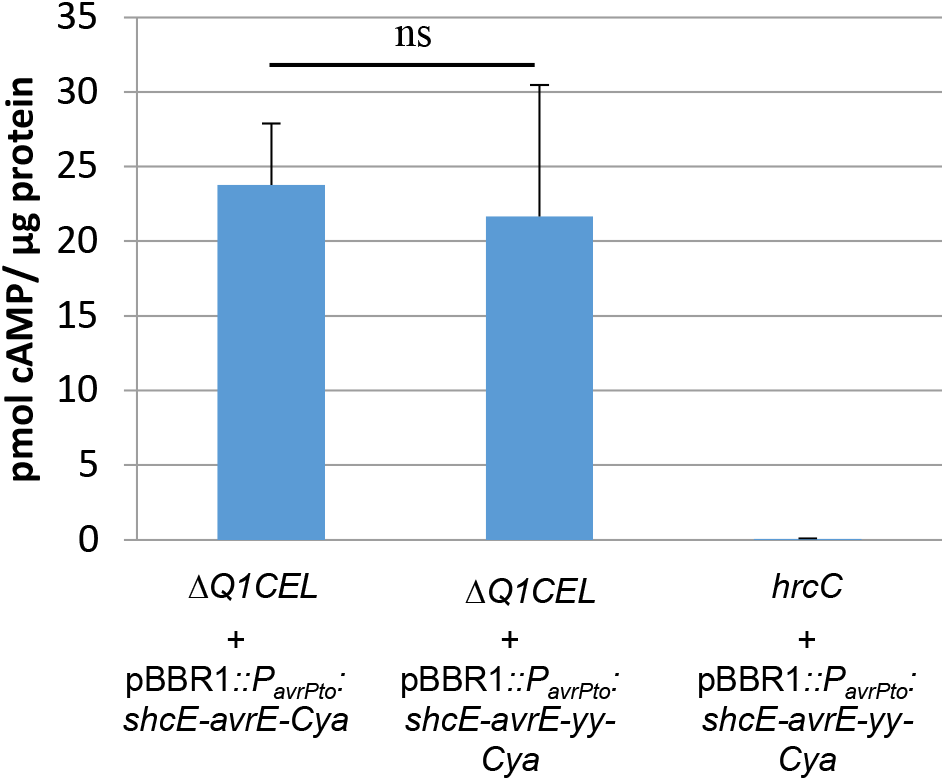
The AvrE-yy mutant protein is translocated by *Pst* DC3000 into plant cells at levels similar to AvrE WT protein in a T3SS-dependent manner. The Cya calmodulin-dependent adenylate cyclase translocation-reporter tag was used. Leaves of 6-week-old *N. benthamiana* were infiltrated with Δ*hopQ1*ΔCEL+pBBR1*:avrF:avrE-WT:Cya*, Δ*hopQ1*ΔCEL+ pBBR1:*avrF:avrE-yy:Cya* or Δ*hopQ1 hrcC+*pBBR1:*avrF:avrE-yy:Cya* at 5×10^7^ cfu/ml. Tissue samples from infiltration areas were collected at 7 h post infiltration and pmol cAMP/μg protein concentrations were determined from duplicate technical repeats. Displayed are means and standard deviations of values from three independent experiments. ns, not significant.

